# Adult survival is reduced when endogenous period deviates from 24h in a non-human primate (*Microcebus murinus*), depending on sex and season

**DOI:** 10.1101/2020.06.24.168765

**Authors:** Clara Hozer, Martine Perret, Samuel Pavard, Fabien Pifferi

## Abstract

Circadian rhythms are ubiquitous attributes across living organisms and allow the coordination of internal biological functions with optimal phases of the environment, suggesting a significant adaptive advantage. The endogenous period called *tau* lies close to 24h and is thought to be implicated in individuals’ fitness: according to the circadian resonance theory, fitness is reduced when *tau* gets far to 24h. In this study, we measured the endogenous period of 142 mouse lemurs (*Microcebus. murinus*), and analyzed how it affects their survival. We found different effects according to sex and season. No impact of *tau* on mortality was found in females. However, in males, the deviation of *tau* from 24h substantially increased mortality, particularly during the inactive season (winter). These results, comparable to other observations in mice or drosophila, show that captive gray mouse lemurs enjoy better fitness when their circadian period closely matches the environmental periodicity. In addition to their deep implications in health and aging research, these results raise further ecological and evolutionary issues regarding the relationships between fitness and circadian clock.

## Introduction

The circadian clock is a pervasive feature, expressing biological rhythms that control a wide range of physiological, metabolic and behavioral traits (Barclay et al., 2012; Kalsbeek et al., 2014; Karatsoreos et al., 2011; Kyriacou and Hastings, 2010; Reinke and Asher, 2016; Richards and Gumz, 2013; Scheer et al., 2013; Wright et al., 2012). It helps coordinating intrinsic biological processes with optimal phases of the daily fluctuating environment. The endogenous period, also called the free-running period or *tau*, represents the duration of a complete circadian cycle (Aschoff, 1960). It lies around 24h in most organisms but expresses variance among individuals (Pittendrigh and Daan, 1976a). It is intrinsically maintained by indirect feedback loops controlling a set of clock genes within the suprachiasmatic nuclei (SCN) (Duong et al., 2011; Gekakis et al., 1998; Lande-Diner et al., 2013; Yu et al., 2002). Every day, the Earth rotation imposes the entrainment of the circadian clock to the 24h light-dark cycles of the environment. Consequently, this daily synchronization between environment and circadian clock is greater in organisms whose endogenous period goes far from 24h and may generate some marginal metabolic or physiological costs that could accelerate aging process and affect survival in the long term.

In that regard, the circadian resonance theory assumes a relationship between *tau* and fitness, via the potential above-mentioned daily metabolic costs: the greater the deviation of *tau* from 24h, the lower survival (Pittendrigh and Minis, 1972). Indeed, drosophila reared under photoperiodic regimens far from 24h displayed reduced survival (Pittendrigh and Minis, 1972; von Saint Paul and Aschoff, 1978). Wyse et al. (2010) found negative correlations between maximum lifespans and *tau* in several species of rodents and primates (Wyse et al., 2010); in a study by Libert et al. (2012), mice with endogenous period close to 24h lived about 20% longer than those with shorter or longer *tau* (Libert et al., 2012). These studies provide evidence that keeping an endogenous period far from 24h increases mortality. We thus wondered if a correlation between *tau* and mortality could be verified in one single primate species.

To address this issue, we focused on an emerging model in neurosciences, the gray mouse lemur (*Microcebus murinus*). This small nocturnal primate originates from Madagascar and displays aged-related impairments similar to those found in humans (Bons et al., 2006; Joly et al., 2014; Languille et al., 2012; Picq et al., 2015), including circadian rhythms alteration, such as locomotor activity fragmentation or sleep deterioration (Aujard et al., 2006; Hozer et al., 2019; Pifferi et al., 2012, 2011). In captivity, the gray mouse lemur can live to age 12 (Languille et al., 2012) and its half-life is approximately 5-6 years (Perret, 1997; Pifferi et al., 2018), whereas the lifespan is significantly lower in the wild (Lutermann et al., 2006). In natural habitat, substantial seasonal variations compel drastic changes in mouse lemur’s metabolism and behavior. The hot rainy season (or summer-like season), characterized by long photoperiod and abundance of resources corresponds to high levels of metabolic activity, as well as reproductive behavior. Rather, during the cooler dry season (or winter-like season), food scarcity and low temperatures force the gray mouse lemur to considerably slow down its metabolism inducing a global fattening and frequent daily phases of hypometabolism (Génin and Perret, 2003; Schmid and Speakman, 2000; Vuarin et al., 2014). These seasonal phenotypic changes are only triggered by the photoperiod: when exposed to photoperiod shorter than 12h, the gray mouse lemurs exhibits a winter phenotype, whereas it displays a summer phenotype when exposed to a photoperiod greater than 12h (Languille et al., 2012; Perret and Aujard, 2001a).

Recently, we reported that mouse lemurs raised under light-dark cycles of 26h exhibited higher daily body temperature and metabolic rate than animals kept in natural lighting conditions (24h), demonstrating the existence of potential metabolic and physiological costs of clock synchronization when endogenous and external rhythms deviate (Hozer and Pifferi, 2020). Do these costs affect the survival of adult individuals? We analyzed how *tau* affected survival in 142 mouse lemurs. In males, the deviation of *tau* from 24h substantially increased mortality, particularly during the inactive season (winter), whereas it did not affect mortality in females.

## Material and Methods

### Animals and ethical statement

All data integrated in the study were taken from the database of the mouse lemur colony of Brunoy (MECADEV, MNHN/CNRS, IBISA Platform, agreement F 91.114.1, DDPP, Essonne, France). These data have been collected between 1996 and 2013. Animals were raised in the laboratory under identical nutritional and social conditions. They were all treated using the same experimental methodology, as described below.

### Housing conditions

All animals were housed in cages equipped with wood branches for climbing activities as well as wooden sleeping boxes mimicking the natural sleeping sites of mouse lemurs, *i.e.* tree holes or cavities. The temperature and the humidity of the rooms were maintained at 25 to 27°C and at 55% to 65%, respectively. In captivity, the artificial daily light cycle within the housing rooms is changed to mimic season alternation, with periods of 6 months of summer-like long days (14h of light and 10h of darkness, denoted 14:10) and 6 months of winter-like short days (10h of light and 14h of darkness, denoted 10:14). Animals were fed *ad libitum* (with fresh fruits and a homemade mixture, see Dal-pan et al., 2011 (Dal-pan et al., 2011) for details).

### Telemetry implants

Recording of locomotor activity (LA) was obtained by telemetry at a constant ambient temperature of 25°C. A small telemetric transmitter weighing 2.5 g (model TA10TA-F10, DataScience Co. Ltd, Minnesota, USA) was implanted into the visceral cavity under ketamine anesthesia (Imalgene, 100mg/kg ip). After surgery, animals returned to their home cage and were allowed to recover for 15 days before start of experiment and continuous recordings of LA. Total recovery was checked by visual inspection of the complete healing of the surgical incision. A receiver was positioned in the cage. Locomotor activity was continuously recorded by the receiver plate, which detected vertical and horizontal movements (coordinate system, DataquestLab Prov.3.0, DataScience Co.Ltd, Minnesota, USA).

### Endogenous period measurement

In order to measure the endogenous period *tau*, individuals included in the survey were isolated and submitted to free-running conditions *i.e.* total darkness during 7 to 30 days. Feeding and weighing were planned at random times of the day, so that no bias could be introduced by human activity in the animals’ facility. After circadian activity measurements, the light regimen was returned to 10:14 light-dark cycles. Endogenous periods were assessed using Clocklab software (Actimetrics, Evanston, IL, USA).

### Mortality data

In total, 142 animals, with 91 males and 51 females were integrated in the study. Thirty-six of them were right-censored (Table 1) because they were alive at the end of the study (*i.e.* 23 march 2020), had been transferred to another laboratory or had been euthanized during experiments requiring sacrifice. The remaining 106 died natural deaths. Dates of birth and natural death or censoring of each individual as well as the dates of season change were known.

**Table 1:** Mean age at death, age at *tau* measurement, number of natural deaths and censored individuals in male and female mouse lemurs. Data are presented as mean ± SD.

|  | Males | Females |
| --- | --- | --- |
| Mean age at death (in years) | 5.98 ± 2.35 | 5.69 ± 2.03 |
| Mean age at *tau* measurement (in years) | 3.10 ± 1.85 | 3.81 ± 2.20 |
| Death from natural causes | 66 | 40 |
| Censored | 25 | 11 |

### Statistical analyses

Because complex interactions exist between sex, season and mortality (Landes et al., 2017; Languille et al., 2012), we treated males and females separately (for an analysis incorporating males and females together, see supplementary materials Table S1 and Figure S1). We first assessed if there was a relationship between *tau* and the age at *tau* measurement, using a Pearson correlation test. For each individual, we then calculated a “Dev.tau” value, which is the absolute deviation of the individual’s endogenous period from 24h (*Dev. tau* = |24 − *tau*|). The individuals entered the study at *tau* measurement, and were left truncated at this age. We then investigated the influence of Dev.tau on survival using multi-variate Cox proportional hazard survival analysis (Cox and Oakes, 1984). The hazard function (defined as the instantaneous risk that the event of interest, *i.e.* death, happens) links a baseline hazard function and a vector of covariables via the following function: *h*(*t*|*y*) = *h*_0_(*t*). *e*^*βy*^, with *h*_*0*_(*t*) being the baseline hazard function of unspecified distribution, *t* being the age, and *β* being the parameter quantifying the impact (effect size) of covariate *y*. We first treated Dev.tau as a continuous variable and we included other additional covariates, that are known to affect mortality in this species: season (integrated as a time-varying covariate), and body mass at *tau* measurement (Kraus et al., 2008; Languille et al., 2012; Pifferi et al., 2018). We also included the following adjustment variables: lineage entered as fixed effect, as well as birth year, year at *tau* measurement, identities of individuals and their mothers entered as random variables. We considered all interactions two by two between fixed parameters. We selected the best models using a backward procedure by calculating second-order Akaike’s Information Criterion (AICc), conserving only models with delta(AICc) <2. Since the use of AICc is not appropriate to predict significance of random effects, we compared the effect sizes of models incorporating the random variables *vs* models that did not, to potentially adjust for interindividual heterogeneity, maternal effects or cohort effects. Verification of the proportionality and the linearity of the models was made using Shoenfeld and martingale residuals. Finally, we constructed parametric Accelerated Failure Time (AFT) models equivalent to the selected Cox models in term of incorporated covariates, to test whether Dev.tau accelerate the speed at which mortality increase with age. We compared them to equivalent parametric Proportional Hazard (PH) models. Both methods assume a Gompertz baseline distribution. This procedure allowed to test whether Dev.tau affected mortality patterns by changing the rate of aging.

## Results

### Distribution of *tau* and correlation with age

As found in many nocturnal species (Pittendrigh and Daan, 1976a), the gray mouse lemur clock oscillates with a period of less than 24h (Fig. 1). Only 2 males and 2 females had an endogenous period greater than 24h. Mean endogenous periods were 23.09 ± 0.58h and 23.10 ± 0.64h in males and females respectively. There was no significant correlation between *tau* and age at *tau* measurement neither in males (r=0.13, p=0.20), nor in females (r=-0.22, p=0.12) (Fig. 2). These two results suggest that endogenous period is both specific to individuals and independent of age in this species.

**Figure 1:**
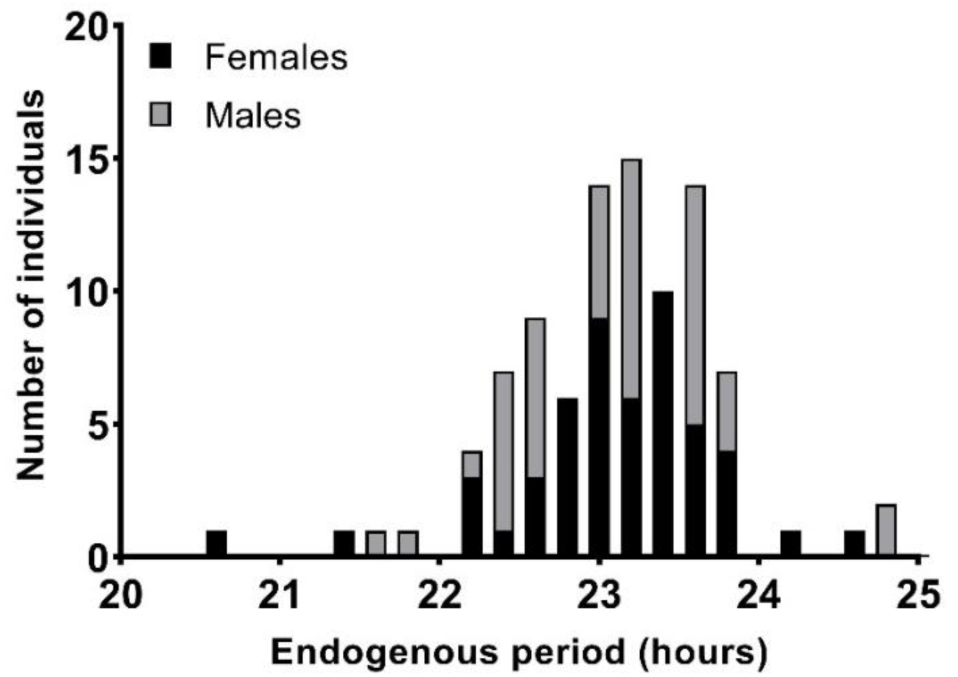
Endogenous period repartition in the 142 mouse lemurs tested. Most of the endogenous periods were less than 24h, only 4 individuals had a *tau* higher than 24h.

**Figure 2:**
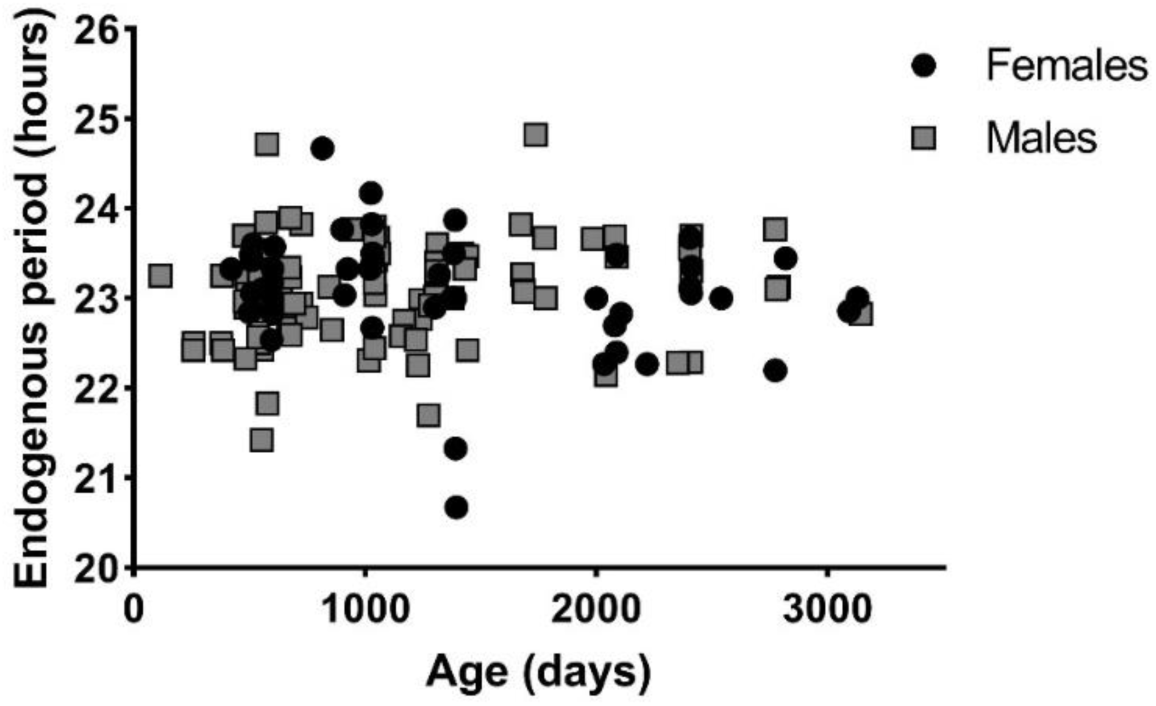
Endogenous periods according to age at measurement.

### The influence of *tau* on survival depends on sex and season

Best selected models for the female and male samples are presented in table 2. They incorporate only variable Dev.tau, season, the interaction between Dev.tau and season and body mass. All models incorporating the variable lineage have larger AICc such that delta(AICc)>2. Individual, maternal and cohort effects did not change the effect sizes of the models. Schoenfeld residuals show that the selected models satisfy the proportionality assumption and martingale residuals did not reveal departure from a linear effect of Dev.tau on mortality. As the effect of Dev.tau remains unchanged among the different selected models and is not modifies by other covariates than season, we chose to treat Dev.tau as a categorical variable into a Kaplan-Meier analysis in order to allow graphic representation. We split each samples (males and females) into three categories, corresponding to tertiles of Dev.tau values (such as every category included an equal number of individuals greater than 15). For females, the tertiles were 0-40min (17 individuals), 40min-1h (17 individuals) and ≥1h (17 individuals). For males, tertiles values were 0-42min (31 individuals), 42min-1h04 (30 individuals) and ≥1h04 (30 individuals) (Fig. 2).

**Table 2:** Estimated effect sizes (± SD) and p-values for all variables retained in the selected models in females, males and males in winter-like.

| Sex | Dev.tau |  | Season |  | Dev.tau:Season |  | Body mass |  | AICc |
| --- | --- | --- | --- | --- | --- | --- | --- | --- | --- |
| | $\beta$ | p | $\beta$ | p | $\beta$ | p | $\beta$ | p | |
| Females | -0.17 ± 0.40 | 0.67 | - | - | - | - | - | - | 196.7 |
| | -1.28 ± 0.95 | 0.18 | -0.49 ± 1.15 | 0.67 | 1.37 ± 1.07 | 0.20 | 0.01 ± 0.01 | 0.11 | 196.5 |
| | -1.12 ± 0.96 | 0.24 | -0.32 ± 1.15 | 0.78 | 1.26 ± 1.08 | 0.24 | - | - | 196.5 |
| | -0.27 ± 0.40 | 0.49 | - | - | - | - | 0.01 ± 0.01 | 0.12 | 196.5 |
| | -0.21 ± 0.41 | 0.60 | 0.88 ± 0.52 | 0.09 | - | - | 0.01 ± 0.01 | 0.13 | 195.7 |
| | -0.12 ± 0.40 | 0.76 | 0.91 ± 0.52 | 0.08 | - | - | - | - | 195.6 |
| Males | 0.99 ± 0.39 | **0.01\*** | 1.45 ± 0.59 | **0.01\*** | -1.38 ± 0.53 | **0.009\*\*** | 0.01 ± 0.01 | 0.45 | 427.3 |
| | 1.04 ± 0.39 | **0.007\*\*** | 1.46 ± 0.59 | **0.01\*** | -1.39 ± 0.53 | **0.008\*\*** | - | - | 425.6 |
| Males in winter-like | 1.17 ± 0.42 | **0.005\*\*** | - | - | - | - | - | - | 144.5 |
| | 1.09 ± 0.42 | **0.009\*\*** | - | - | - | - | 0.02 ± 0.01 | 0.09 | 144.2 |

Dev.tau did not influence survival in females (Table 2 and Fig. 2A).

In males, however, Dev.tau affected negatively and significantly the survival. Our best model predicted that every supplementary hour of Dev.tau (*i.e.* every supplementary hour of *tau* deviation from 24h) multiplied the risk of death by 2.82 (=*e*^1.04^, Table 2). Dividing the sample into tertiles of Dev.tau, median survival ages were 2637 days (7.22 years), 2170 days (5.94 years) and 1704 days (4.67 years) in the three represented groups respectively, *i.e.* a reduction of 35% of median survival between the two extreme curves. Maximum longevities were 3363 days (9.21 years), 3772 days (10.33 years), and 3915 days (10.73 years) in the three groups respectively (Fig. 2B).

Besides, the interaction between Dev.tau and season in males was significant as well (Table 2). When only considering the winter-like season, the negative effect of Dev.tau in males on survival was amplified: every supplementary hour of Dev.tau multiplied the risk of death by 3.22 (=*e*^1.17^, Table 2). Dividing the sample into tertiles of Dev.tau, median survival ages were 3180 days (8.71 years), 2844 days (7.79 years) and 1413 days (3.87 years) in the three represented groups respectively, *i.e.* a reduction of 56% of median survival between the two extreme curves. Maximum longevities were 3363 days (9.21 years), 3772 days 10.33 years), and 3871 days (10.61 years) in the three groups respectively (Fig. 2C).

Parametric AFT models were equally but not better supported by the data than proportional hazard model (data not shown).

**Figure 2:**
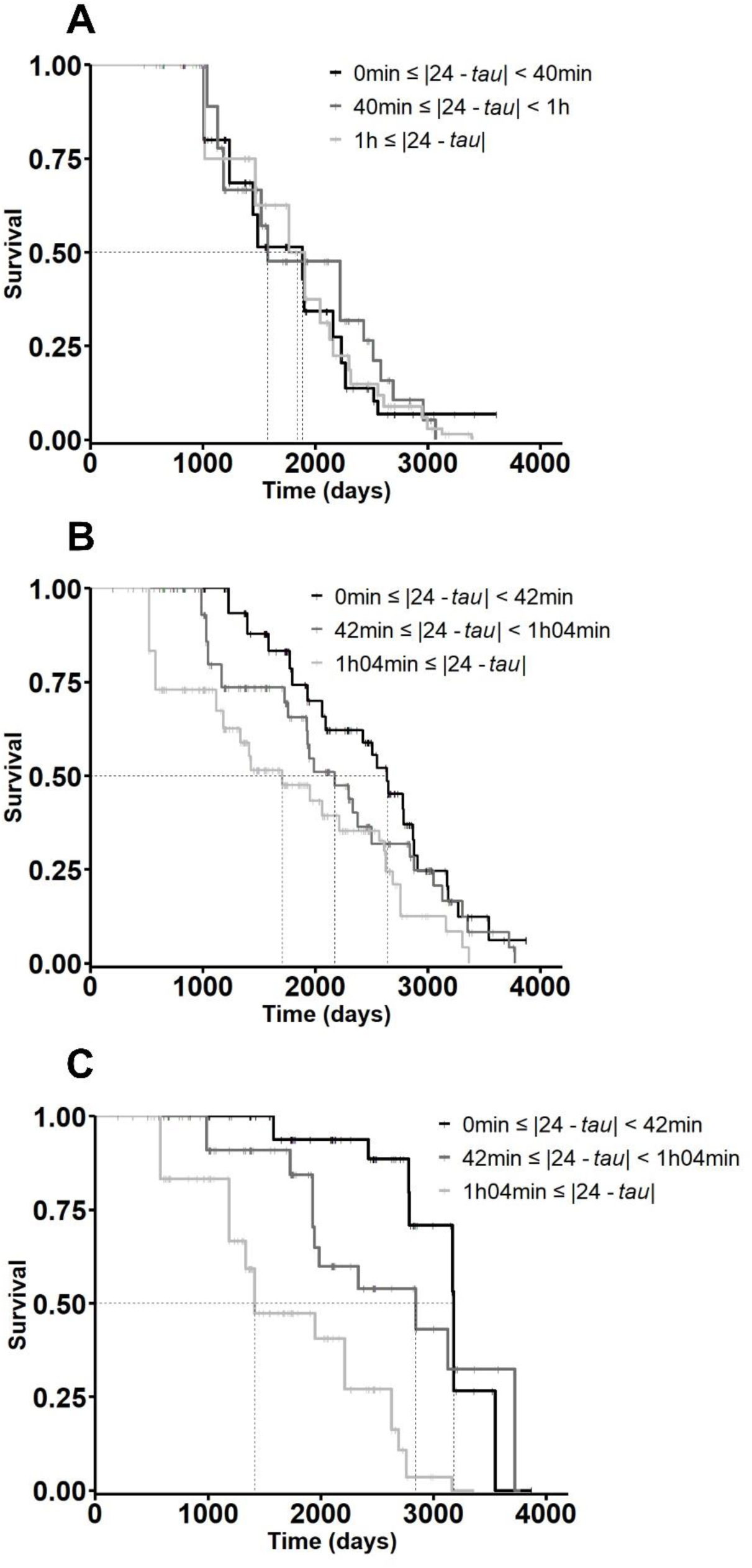
Effect of Dev.tau on the survival of females (A), males (B) and males in winter-like season (C) with increasing age. Individuals were distributed in tertiles to allow graphic representation, corresponding to specific cut-off of absolute deviation of *tau* from 24h. Dotted lines correspond to median survivals. Small solid vertical bars correspond to censored data.

## Discussion

This study aimed at exploring the relationship between the endogenous period *tau* and survival in a non-human primate. Our results show no impact of endogenous period on survival in female mouse lemurs. On the other hand, male individuals with endogenous period close to 24h were those experiencing a better survival. This effect was particularly significant in winter-like season, corresponding to the non-mating and inactive season. Individuals with *tau* far from 24h had a reduction of median survival up to 56% compared to individuals with closer *tau*, suggesting a high adaptive value of maintaining an endogenous period close to 24 h. It is still not clear, however, why *tau* exhibits such important inter-individual variance but the deviation of *tau* from 24h may bring some advantage in seasonal adaptation by stabilizing the phase angle (Pittendrigh and Daan, 1976b). So far, the relationship between endogenous period and survival had been assessed in fruit flies and mice (Libert et al., 2012; Pittendrigh and Minis, 1972). In natural conditions, it was also reported that mice with a mutation in the enzyme casein kinase 1ε exhibited a short *tau* (∼22h) and displayed lower survival and reproduction performances when released in outdoor enclosures (Spoelstra et al., 2016). Only one study found a similar relationship in primates, but it applied a totally different experimental protocol, based on interspecific comparisons and maximum lifespans taken from the literature (Wyse et al., 2010). To our knowledge, our results are the only ones to report an influence of *tau* on survival in a single primate species.

Why are the endogenous period and survival related to each other? Pittendrigh and Minis suggested that the daily resetting of the biological clock onto the 24h of the environment would engender daily marginal metabolic costs that would accelerate aging and affect survival: the impact of these costs on longevity would be proportional to the deviation of *tau* from 24h. In that respect, the median survival ages found in our experiment show clearly that the more *tau* gets far from 24h, the more the survival is low. Recently, we observed that mouse lemurs kept in light-dark cycles far from their endogenous period (26h) exhibited higher resting body temperature and higher energy expenditure, associated with lower cognitive performances (Hozer and Pifferi, 2020). This study suggests that living under photoperiodic regimen far from endogenous rhythms leads to physiological, metabolic and cognitive costs for the organism. It could partially explain the link that have been made between *tau*, longevity and aging. Indeed, biological clock and aging processes are assumed to influence one another. Aging is often associated with rhythm fragmentation, phase advance, sleep disorders, decrease of rhythm amplitude, SCN atrophy (Engelberth et al., 2014; Jazwinski et al., 2017; Monk, 2005; Nakamura et al., 2011; Zhdanova et al., 2011). Besides, the deterioration of the biological clock is alleged to accelerate and stand at the heart of aging mechanisms (Banks et al., 2016; Kondratov et al., 2006; Yang et al., 2016; Yu and Weaver, 2011). In this context, the endogenous period could modulate the rate of aging and indirectly influence organisms’ mortality. For example, it was reported that plasma level of interferon-γ, a well-known biomarker of aging, was negatively correlated to survival in gray mouse lemurs. Interestingly, the plasma level of interferon-γ was also negatively correlated with the endogenous period, *i.e.* the individuals with *tau* close to 24h displayed lower aging biomarkers and greater survival (Cayetanot et al., 2009). These observations, along with our results, underline the relevance of an acute and coordinated circadian pacemaker, including endogenous rhythms resonating with the environment cycles, to enhance survival. In nature, one may wonder if the endogenous period substantially impacts fitness, since individuals often die before aging. An interesting study would be to place wild type individuals in semi natural conditions, and to monitor their survival depending on their endogenous period. Another interesting perspective could be to investigate the relationship between *tau* and other aspects of fitness, such as reproductive events or juveniles’ recruitment.

Why does the endogenous period influence survival in males but not in females? Libert’s study focused exclusively on males, but curiously, Pittendrigh and Minis found similar results on males and females, which is intriguing, regarding our results, since insect and mammal clocks are known to display significant similarities (Helfrich-Förster, 2004; Stanewsky, 2003). We have no clear explanation, but it should not be forgotten that the study included more males than females, increasing the risk of type II statistical error in females. Otherwise, this difference between sexes may be related to mammal sexual circadian specificities, even though a bias towards research on male’s circadian clock makes exhaustive comparisons between sexes difficult (Krizo and Mintz, 2015). Some studies report though that the influence of internal desynchronization on sleep-wake disorders is considerably greater in females than in males, some other mentioned a slower synchronization in females (Bailey and Silver, 2014; Duffy et al., 2011; Goel and Lee, 1995; Wever, 1984). The daily costs of resynchronization due to *tau* deviation may be negligible compared to more important circadian alterations. Unfortunately, sexual comparisons in circadian parameters are little studied in the gray mouse lemur. However it is known that female mouse lemurs deal with seasonal transitions and metabolic costs differently than males, particularly during aging (Perret and Aujard, 2001a). Facing an environmental metabolic stress, females mouse lemurs seem to better manage their energy expenditure and exhibit more flexible physiological response (Génin and Perret, 2000; Noiret et al., 2020; Schmid, 1999; Terrien et al., 2010). During aging, males body weight variation collapse whereas females keep displaying clear seasonal mass variations, suggesting a better management of resources (Perret, 1997; Perret and Aujard, 2005). Therefore, females may better deal with generated daily metabolic costs due to circadian clock reset and thus would not display survival impairments.

Why does *tau* influence survival in winter-like more than in summer-like? The explanation could lie in the huge physiological and metabolic changes experienced by the gray mouse lemurs between winter-like and summer-like photoperiods. The species has the particularity of being heterothermic, allowing daily phases of hypometabolism (torpor) that occur almost exclusively during the winter-like season, in order to cope with environmental food scarcity and to save substantial amount of energy, even in captive conditions (Aujard et al., 1998; Perret, 1998). It has been shown that circadian clock and torpor use are closely related (Perret and Aujard, 2001b). For example, light pollution significantly modifies circadian expressions, with negative repercussions on torpor (Le Tallec et al., 2013). If the circadian clock oscillates too far from environmental periodicity, it may imply affectations of torpor efficiency and be deleterious in the long run. Inversely, in summer-like season, during which torpor hardly ever happens, the activation of reproduction investments may require so much energy that reproduction investments in males could overwhelm daily costs engendered by clock synchronization. In addition, IGF-1 (Insulin-like Growth Factor 1) rates exhibit an age-related decrease in male mouse lemurs only during the winter-like season, whereas they remain high and stable during the summer-like season. Interestingly, the winter-like IGF-1 rates are also predictive of survival (Aujard et al., 2010). It could thus be worthwhile to combine data from *tau* and IGF-1 in this species, as it is known that circadian rhythms and IGF-1 are closely related (Breit et al., 2018; Crosby et al., 2019).

In this study, we showed that Dev.tau was not influenced by age between individuals, but we made the hypothesis that *tau* was constant over each individual lifetime. It is however not clear if time and more specifically aging do influence the endogenous period or not. In several mammalians, even within the same species, increasing, decreasing or constant endogenous periods with age have been found (Davis and Viswanathan, 1992; Kendall et al., 2001; Monk and Moline, 1989; Pittendrigh and Daan, 1976a; Possidente et al., 1995). In the gray mouse lemur, the same conflicting observations lead to suggest that *tau* evolution during aging is due to epigenetic variations between individuals and underline once again that aging is an individual-dependent phenomenon (Aujard et al., 2006; Hozer et al., 2019; Schilling et al., 2001). A longitudinal approach is necessary to bring further exhaustive information on the relationship between *tau* and aging.

To conclude, our results highlighted the negative impact of great deviation of *tau* from 24h on survival in adult captive male mouse lemurs, even if further investigations are needed to elucidate the ecological implications of sex and season dependence. Our findings show that resonating circadian clocks are obviously highly adaptive features of living organisms and underline the importance of correct clock resetting to enjoy better health. They may have some interesting applications in human societies, where light pollution and drifted ways of life modulate circadian clock adjustments to external world.

## Acknowledgments

We are grateful to Aude Anzeray and Isabelle Hardy for their participation in the data collection.

## Author Contributions

Conceptualization, CH, FP and MP. Investigation, CH and MP. Methodology and Formal Analysis, CH, SP and FP, Writing-Original Draft, CH; Writing-Review & Editing, CH, MP, SP and FP. Supervision, FP.

## Materials and Correspondence

Correspondence and material requests should be addressed to Fabien Pifferi.

